# Prediction of KIR3DL1/Human Leukocyte Antigen binding

**DOI:** 10.1101/2024.05.03.592082

**Authors:** Martin Maiers, Yoram Louzoun, Philip Pymm, Julian P. Vivian, Jamie Rossjohn, Andrew G Brooks, Philippa M. Saunders

## Abstract

KIR3DL1 is a polymorphic inhibitory Natural Killer (NK) cell receptor that recognizes Human Leukocyte Antigen (HLA) class I allotypes that contain the Bw4 motif. Structural analyses have shown that in addition to residues 77-83 that span the Bw4 motif, polymorphism at other sites throughout the HLA molecule can influence the interaction with KIR3DL1. Given the extensive polymorphism of both KIR3DL1 and HLA class I, we built a machine learning prediction model to describe the influence of allotypic variation on the binding of KIR3DL1 to HLA class I. Nine KIR3DL1 tetramers were screened for reactivity against a panel of HLA class I molecules which revealed different patterns of specificity for each KIR3DL1 allotype. Separate models were trained for each of KIR3DL1 allotypes based on the full amino sequence of exons 2 and 3 encoding the *α*1 and *α*2 domains of the class I HLA allotypes, the set of polymorphic positions that span the Bw4 motif, or the positions that encode *α*1 and *α*2 but exclude the connecting loops. The Multi-Label-Vector-Optimization (MLVO) model trained on all alpha helix positions performed best with AUC scores ranging from 0.74 to 0.974 for the 9 KIR3DL1 allotype models. We show that a binary division into binder and non-binder is not precise, and that intermediate levels exist. Using the same models, within the binder group, high- and low-binder categories can also be predicted, the regions in HLA affecting the high vs low binder being completely distinct from the classical Bw4 motif. We further show that these positions affect binding affinity in a nonadditive way and induce deviations from linear models used to predict interaction strength. We propose that this approach should be used in lieu of simpler binding models based on a single HLA motif.

## 1 Introduction

Natural killer (NK) cells play a key innate role in innate surveillance of infected or transformed cells^1^. Their activation is governed by the balance of signals received from a diverse array of inhibitory and activating receptors. Perhaps the best described inhibitory receptors belong to the Killer cell Immunoglobulin-like Receptor (KIR) family which recognize allotypic subsets of HLA class I molecules (HLA-I). These interactions are important during NK cell development driving the acquisition of functional potential through a process termed education and additionally facilitate NK cell recognition of target cells which lack self-encoded HLA-I as can occur following infection, transformation, or allogenic transplantation.

The KIR genes, which are located on chromosome 19q13 exhibit remarkable genomic diversity. KIR gene content varies between individuals who can possess from 7 to 14 activating and inhibitory KIR genes^2,3^. Furthermore, in addition to significant haplotypic differences in the *KIR* gene content, there is significant polymorphism within individual genes with 2,219 alleles described to date in the Immuno-Polymorphism Database^4^ Release 2.12 (2023-12).

The impact of polymorphism within individual KIR genes is best understood in the context of *KIR3DL1* that encodes an inhibitory receptor specific for HLA-I molecules that possess a serological motif termed Bw4, a polymorphic region that spans residues 77–83 on the α1-helix of the HLA-I molecule. Early studies^5^ suggested that Bw4 allotypes that possessed Ile at position 80 preferentially inhibited NK cell activation, although certain allotypes such as HLA-A*25:01 that possessed Ile80 were not well recognized implying that the docking site for KIR3DL1 on HLA-I was not simply defined by the motif. The crystal structure of the KIR3DL1*001 bound to HLA-B*57:01 ^6^ revealed that in addition to residues within the Bw4 motif, KIR3DL1 bound two additional regions that are highly conserved across HLA-I allotypes. Moreover, in addition to residues on the heavy chain itself, KIR3DL1 also made direct contacts with the HLA-I-associated peptide. Thus, polymorphism within and outside the Bw4 motif could impact the interaction with KIR3DL1 either by directly shaping the microarchitecture of both the α1 and α2 helices or indirectly by dictating the sequence and conformation of the peptides bound to the HLA^6–10^.

Our understanding of the HLA-Bw4 preferences of KIR3DL1 is further confounded by polymorphism within the *KIR3DL1* locus. To date, 184 allelic variants of the *KIR3DL1* gene have been described encoding 92 distinct amino acid sequences. Phylogenetically these allotypes span three main lineages based on sequence differences across the three extracellular domains (D0–D1– D2) of the KIR3DL1 molecule^4,11,12^. There are two diverse inhibitory lineages comprising *KIR3DL1*005*-like and *\*015*-like alleles, and a third lineage which is much more constrained at a population level and consists primarily of the activating *KIR3DS1*013* allele^13^. *KIR3DL1* polymorphism was initially associated with differences in cell surface expression of KIR, with allotypes such as KIR3DL1*001, *002, and *008 expressed at high levels, *005 and *009 at lower levels, and *004 being largely retained intracellularly^14,15^. However this polymorphism may also directly affect recognition of HLA class I with polymorphisms at 238 and 283 associated with differences in recognition of HLA-Bw4 ligands^7,16–18^.

This complex interaction between allotypic variants of both HLA-Bw4 and KIR3DL1 was demonstrated in binding analyses using soluble forms of KIR3DL1 to bind an array of HLA-I-coated beads. These data revealed distinct hierarchies for different KIR3DL1 allotypes with the KIR3DL1*005 having a broader range of ligands than allotypes such as KIR3DL1*015^8,19^. The data also confirmed that while HLA-I allotypes possessing Ile80 are typically amongst the strongest binding HLA-I allotypes, Ile80 was also present in allotypes that bound weakly to KIR3DL1 providing further support for the notion that polymorphism outside of the Bw4 motif is responsible for the interaction with KIR3DL1.

Given the extensive polymorphism of both the KIR3DL1 and HLA class I, it is not practical to experimentally test the binding of each KIR3DL1-HLA pair. Lacking a clear amino acid-based rule for the binding of specific KIR3DL1 and HLA class I allotypes, we here train and evaluate a continuous model for the strength of such binding.

## 2 Material and Methods

### Tetramer experiments

KIR3DL1 tetramers were screened for reactivity against a panel of HLA-I molecules which revealed different patterns of specificity for each KIR3DL1 allotype. HLA-I recognition by KIR3DL1 allotypes was assessed through binding of KIR3DL1 tetramers to beads coated with a panel of 100 different HLA-A, -B, and -C molecules (LABScreen HLA Class I Single Antigen; One Lambda). These experiments are described in detail previously^19^.

### KIR binding affinity measurements

KIR binding affinity values were determined over a series of 3 runs per KIR allotype. The results were averaged and normalized against the maximal response to produce a matrix of raw binding values for each HLA and KIR combination, distributed between 0 and 1.

### 2D-clustering

The log of the raw matrix above was clustered using hierarchical clustering with average link clustering on both the KIR allotypes and the HLA allotypes using a Euclidean distance between samples. The results presented are ordered following the clustering in both directions. For the HLA clustering, each HLA was represented by its nine-dimensional vector of normalized binding affinities to the nine KIR3DL1. The opposite was done for the KIR3DL1 allotype representations.

### Logo plots

Logo plots were computed using a web-tool called two sample logo^20^. Sequences were grouped into three clusters of HLA allotypes based on the hierarchical clustering above. All sequences were then aligned. The high and low binding levels were compared with the non-binder clusters. Then, the low and high binding groups were compared one to each other. The samples were compared using a binomial test. The presented value represents the amino acids with a significant difference at a 0.05 level with a Bonferroni correction^21^.

### Machine learning and regression

The amino acid sequences of the different HLA allotypes were translated into amino acid-position pairs (for example R6 represents an arginine at position 6). Positions for which there was no variability in all studied HLA allotypes were removed from the samples. For example, if position 4 along the HLA always had a serine, it was removed from the analysis.

Each HLA allotype was then represented as a binary vector with 1 in the appropriate position if the HLA had the appropriate amino acid in the appropriate position. For example, an HLA that starts with RA would have values of 1 in positions R1 and A2, and 0 in all other amino acid possibilities for position 1 and 2 (e.g., S1 and L2). This is a one-hot representation with non-polymorphic encodings removed (further denoted as one-hot vectors). We then applied linear or kernel-based predictions on the log of the binding affinity. Zero values were replaced by a minimal value (1% of the minimal positive value to avoid a log of zero values).

The prediction was performed using different possible inputs, either 1) all positions in the second and third exons of the HLA allele, 2) only the six residues associated with the Bw4 motif (defined as [76 77 80:83]), or 3) no loops (defined as [16:19,39:44,49:56,86:90,106:107,128:131,137:140,151:152,176:179,180:183]), residues encoding alpha helices (defined as [57:85,141:175]) or as the exon 2 and 3, without loops or without helices.

When the regression was performed only on the known Bw4 positions, the one-hot vectors themselves were used. When regression was performed using the full sequence, first either a Multiple Correspondence Analysis (MCA) ^22^ or a Principal Component Analysis (PCA) were performed over all the one-hot vectors. Then machine learning was performed on the projection over the first K MCA vectors. In the current application, K was set to be 7, based on the decrease in the contribution to variance beyond the 7^th^ Eigenvector. The results presented for the KIR binding prediction are an average over five cross validations. In all learning tasks, a 5-fold cross validation was performed with the percentage of training and test samples was 80% and 20% respectively, unless explicitly stated otherwise. For each cross-validation, each HLA allele was either always in the test or always in the training set for all KIR3DL1 allotypes. The following classifiers were used:

A. Support Vector Classification (SVM) - The implementation of this method was based on *libsvm* ^23^. The penalty parameter of the error term (the “Box Constraint”) was set to 0.01. The kernels tested were linear and polynomial and a “balanced” class weight was used, implying that the box constraint is normalized to be inversely proportional to the class size.
B. MLVO - Formally, assume samples that have both binary and continuous labels (not necessarily both labels for all samples). We assume that continuous observations are monotonically related to the binary classifications. Each sampled point can have either one of the two label types or both. When both are used the resulting kernel machines based optimization problem has been denoted the Multi-Label-Vector-Optimization^24^.

More complex methods, such as Random Forest, XGBoost and Neural networks had lower performance scores.

The precisions of the classifiers were computed using the AUC (area under the sensitivity-specificity curve).

### Phylogeny

For the comparison between different KIR, the Hamming Distance ^25^ between the sequences of the KIR alleles was computed, using the amino acid sequence over the entire KIR gene, following gene alignment between all the studied KIR.

### Webtool

A webtool (https://kir-hla.math.biu.ac.il/Home) was developed using the python flask framework that implements the predictions from the model based on the amino-acid sequence of the input HLA allele. The output is based directly on the matrix of beta values from 11 models: overall binding, high vs low binding, and KIR allele specific models for the 9 training allotypes: KIR3DL1*001, *002, *004, *005, *008, *009, *015, *020, *029. Supplemental Figures 1, 2 and 4 shows the response of the tool to an input for specific allotypes. Supplemental Figure 3 shows the response of the tool to an input of an HLA Class I amino acid sequence. The webtool ignores AA that are missing (‘*’).

## 3 Results

### Analysis of binding data

Inhibitory KIR share a common binding site on HLA-I spanning residues in both the *α*1 and *α*2 helices. Notably, KIR2DL1/2/3 and KIR3DL1 all bind to residues 145, 146 and 149-151, all of which are highly conserved across HLA-A, -B and -C allotypes^26,27^. The other major KIR docking site spans residues 69-84 which spans both the C1 and C2 serological determinants as well as the determinants of Bw4 and Bw6 epitopes. At positions 80 and 83, there are five common patterns across HLA-I allotypes, two of which correspond to Bw4 allotypes (I80/R83 and T80/R83), K80/G83 and N80/G83 denote C1 and C2 allotypes respectively while T80/G83 characterizes the bulk of HLA-A allotypes (Table 1). Notably, HLA-B molecules that possess the Bw6 motif in lieu of Bw4, also possess the N80/G83 combination present in C2 allotypes.

**Table 1.** Class I common polymorphisms at positions 80 and 83 by epitope name and locus.

| Pos80 | Pos83 | C1/C2 | Bw4 +/- | Loci |
| --- | --- | --- | --- | --- |
| N | G | C1 | - | B C |
| T | G |  | - | A |
| I | R |  | + | A B |
| T | R |  | + | B |
| K | G | C2 | - | C |

To better define how these combinations impact HLA-KIR binding, we have analyzed the binding of 96 common HLA-A, B and C allotypes to nine distinct KIR3DL1 allotypes (Figure 1). As expected, regardless of KIR3DL1 allotype, HLA-I allotypes that possessed either I80/R83 or T80/R83 exhibited superior binding to other combinations. Interestingly, the K80/G83 (C1) and N80/G83 (C2) combinations while on average not binding as well as the Bw4 allotypes, nevertheless showed significant reactivity with KIR3DL1 compared with the T80/G83 allotypes (Mann Whitney test between K80/G83 or N80/G83 and T80/G83 *P*<.001 for all KIR3DL1 allotypes). Additionally, there were significant differences in recognition patterns across individual KIR3DL1 allotypes, with KIR3DL1*005 being the most distinct evidenced by greater capacity to bind N80/G83 allotypes and even a number of T80/G83 allotypes. Nevertheless, HLA allotypes with I80 and T80 (both with R83) were associated with high binding to KIR3DL1. K80 and N80 can be either low-level binders (typically on C allotypes, but some B alleles, mainly to KIR3DL1*005) or non-binders (typically on B allotypes) and T80/G83 do not bind to KIR3DL1, except for KIR3DL1*005. The variation in KIR3DL1 binding within each of these HLA subgroups reinforces the notion that polymorphisms outside of regions 77-83 of the *α*1 helix can significantly impact HLA class I recognition by KIR3DL1.

**Figure 1:**
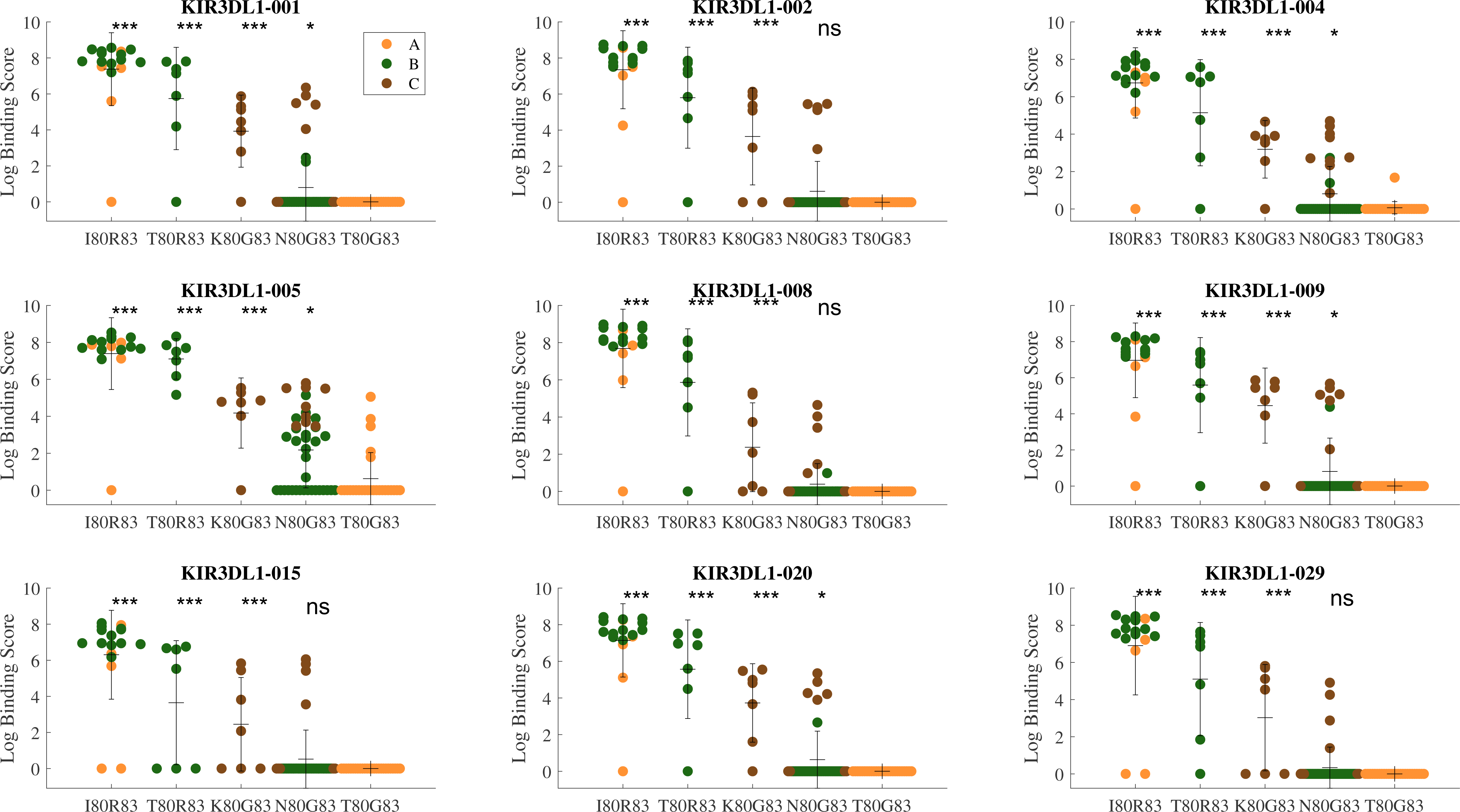
Binding data by KIR3DL1 allotype. The log binding score is plotted for each KIR3DL1 allotype across the HLA allotypes in the panel. Each subplot is a different KIR3DL1 allotype. Within each subplot, HLA are grouped by their amino acid residue combinations at positions 80 and 83 (very rare combinations were ignored). The colors represent the HLA locus. Each group is compared to the lowest binding group (* p<0.05, ** p<0.01,*** p<0.001,ns Non-Significant).

Given the importance of polymorphisms outside the 77-83 region, a computational prediction tool is needed to extend the *in vitro* based binding measurements to other HLA and KIR3DL1 allotypes. Two types of predictions are proposed. A prediction tool for a KIR3DL1 allotype where we have HLA-binding measurements (the nine alleles in Figure 1), but not-covered HLA and a second prediction is required when there are no experimental data for the KIR3DL1 allotype.

To estimate the fraction of the population that require the two predictors, we evaluated the population coverage of the current experimental panel of KIR3DL1 and HLA (Figure 2) and found, based on allele frequencies in the literature, that 6-24% of the of the KIR3DL1 allele frequency distribution was not covered ^11^ and 11-23% of the HLA allele frequency distribution was not covered ^28^. On average, 76% of the HLA-KIR3DL1 allele combinations are covered (green rectangles – Figure 2A). The coverage is maximal in European and Asian populations and is less in African populations as expected due to their great genetic diversity (Figure 2A). Despite this broad coverage for individual KIR3DL1-HLA allotypic combinations, when full HLA class I genotypes across populations were considered (i.e. the combined presence of six HLA class I alleles), the number covered in the current panel reduces to 8% in African populations, and to 34 % in European populations (Figure 2B). Thus, even in the best covered population, two thirds of the population have at least one HLA allele for which a computational prediction is required.

**Figure 2:**
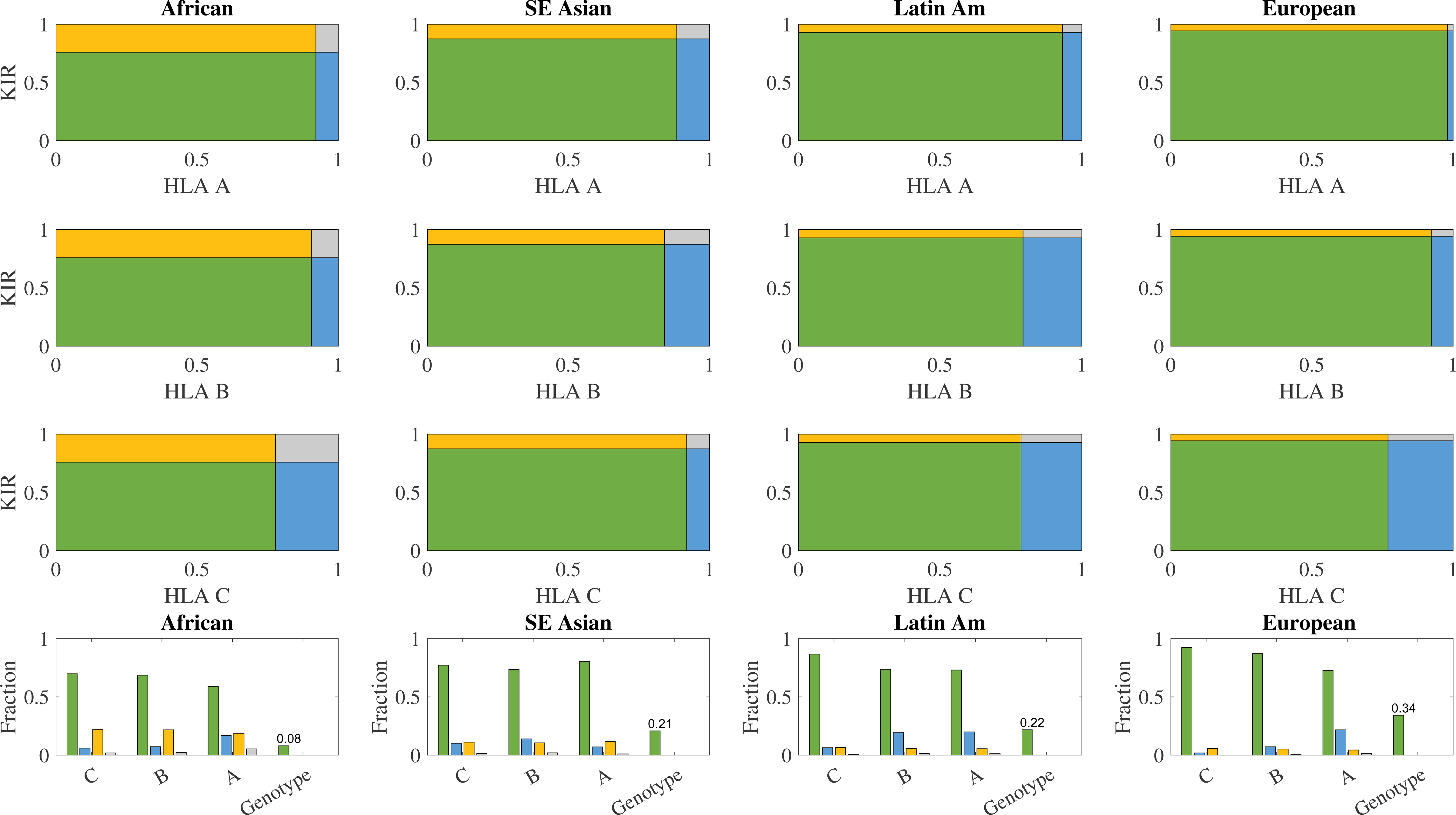
Population Coverage: The binding data can be used to make assessments for particular KIR3DL1 and HLA class I allotypes. However, in order to make this assessment comprehensive (e.g., in clinical applications) prediction is needed to address the gaps in population coverage which can be over 20% and disproportionately affect non-European populations. **A.** The frequency of HLA and KIR3DL1 allotypes among African, Asian, Latin American and European populations was evaluated for HLA-A (**i**), HLA-B (**ii**) and HLA-C (**iii**). Within each box, green regions represent the fraction of the population where both HLA and KIR3DL1 binding information are covered in the current study. Yellow and blue are the presence of either KIR or HLA data respectively and gray is where neither are present in the current dataset. **B.** Summary of the covered fractions along with the fraction of the genotypes composed of two of each HLA allotype fully covered in the experimental binding measures in the current study. For all populations, assuming independence, this fraction is below 0.35.

To better define the residues in HLA-A and -B that were responsible for differences in KIR3DL1 binding we clustered the KIR3DL1 and HLA alleles in the experimental binding data using hierarchical clustering. KIR3DL1 allotypes, *002, *008, *009, *020 and *029 were highly similar, while *005 and *004 were the most divergent (Figure 3A). These differences only partially match the sequence phylogeny with *002, *008, *009, *020 and *029 in the *015-like lineage and *004 and *005 in the *005-like lineage^11^. Clustering of HLA allotypes was performed on all KIR allotypes (i.e. each HLA allotype is represented by its vector of log binding affinities to each KIR allotype). Three clusters of binding were observed based on the top three clusters of average link hierarchical clustering, which we have labeled: high binders, low binders, and non-binders. (Figure 3A).

**Figure 3:**
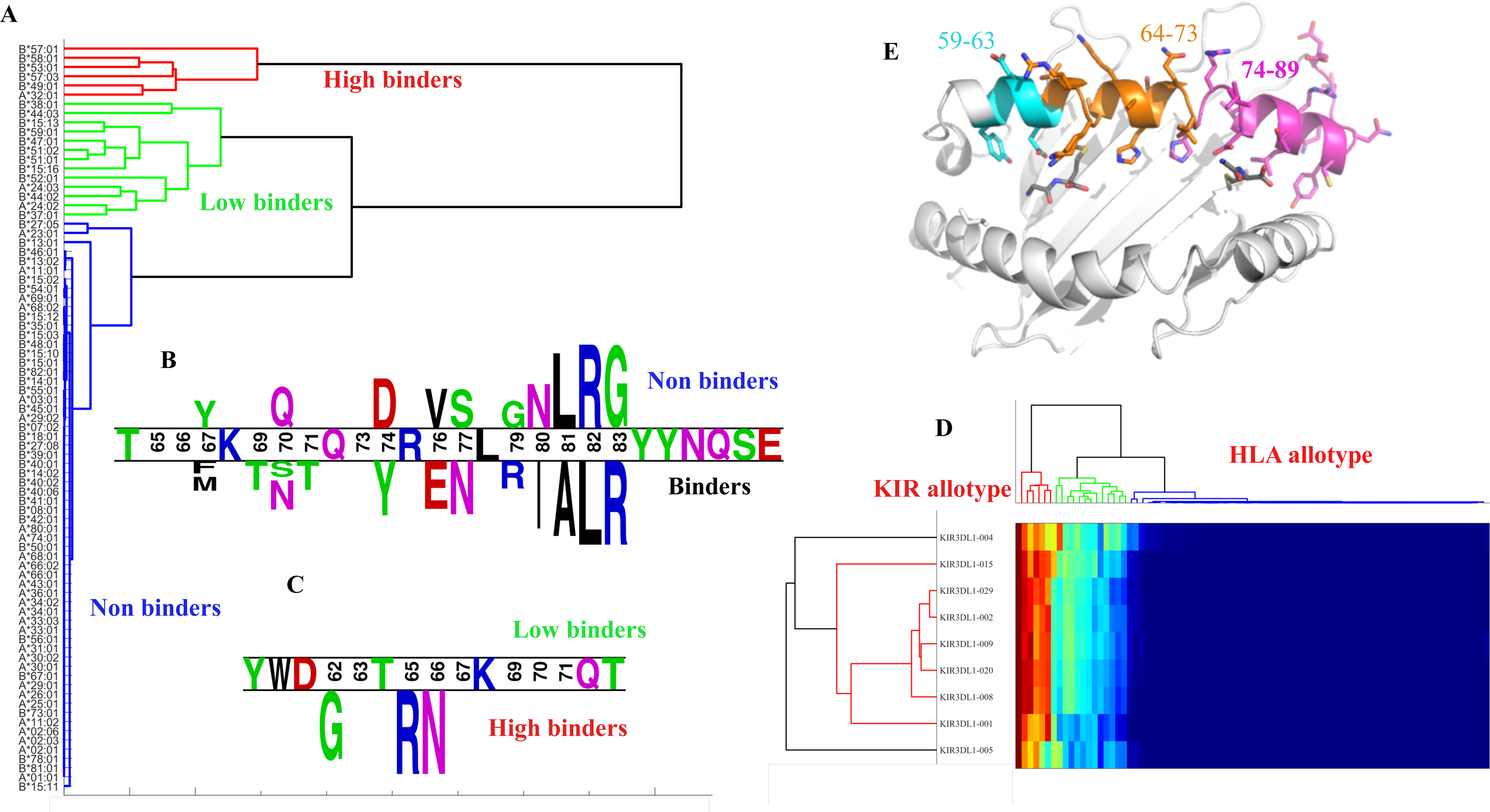
Clustering of binding data and logo plots. **Left plot (A).** An expanded and rotated view of the HLA allotypes from the 2-dimensional clustering to reveal three distinct patterns of binding across all 9 KIR allotypes: high-binder (red), low binders (green) and non-binders (blue). **Lower left plots (B & C)** Logo diagram of the amino acid residues of the HLA-A and HLA-B allotypes in the panel shows that while the Bw4 motif (positions 77-83) has the largest enrichment or depletion for residues associated with binding vs non-binding (both high and low), other positions are significantly enriched or depleted. Below it is a logo diagram generated from a similar comparison of the amino acid sequences of HLA alleles that encoded high vs low binding allotypes. **Lower right plot (D).** 2-dimensional clustering demonstrating relationships between KIR3DL1 allotypes (note the more promiscuous binding of KIR3DL1*005). **Upper right plot (E).** Highlighted are relevant regions on the *α*1 domain of the HLA class I molecule (PDB accession 6TDQ). The purple residues comprise positions 74-89 which were the most significant in the binding prediction. All colored positions are the no-loop regions. Blue residues are positions 59-63, which were significant for the low to high binder predictions, and orange residues are 64-73 that were important for both predictors. Positions not colored showed no significant enrichment or depletion.

We next compared all HLA-I allotypes classified as high or low binders (red and green clusters) to the non-binding cluster (blue cluster) and computed the amino acid residues significantly contributing to the binder vs non-binder split using a two-population logo plot, where the size of each letter is proportional to its contribution to the difference between the populations (Figure 3B-C). Although the contribution to binding is highest at positions 80-83, with a clear IALR (positions 80— 83) signature motif for binders and NLRG motif for non-binders, there are significant contributions from positions extending back to residue 67, which may be due to linkage disequilibrium. Interestingly, not all binders have the full IALR signature (such as HLA-B*44:02 which has a Thr at position 80), but each of these residues contributes significantly to the difference between binders and non-binders. Importantly, all amino acids within the *α*1 and *α*2 domains were considered in the analysis, however only residues with variation substantial enough to produce a signal were included in the model further developed below. Residues that were broadly predictive of binding strength were only found within the *α*1 domain but this does not preclude the impact of unique polymorphisms on the *α*2 helix on specific allotypes such as positions 145 for HLA-B*13:01 and 149 for HLA-A*25:01 that have been reported to impact KIR3DL1 recognition^10^. To further detect the residues associated with the distinction between low and high binders, we performed a similar analysis focusing only on the difference between the high and low binders (Figure 3D). The amino acid residues that significantly contributed to this distinction were at positions 62, 65 and 66, all of which form part of the A pocket which accommodates peptide residues (Figure 3E).

These highlighted positions compose together most of the outward pointing amino acids in the alpha 1 helix, with the classical Bw4 motif at the C-terminal peptide end of the HLA-I. Regions further from the C-terminal peptide have a lesser effect on the KIR3DL1 binding reflected by the decreasing size of the letters in the logo plot from right to left (Figure 3C).

### Binding prediction

Given the extent of allelic diversity in both the *HLA-A, -B* loci and *KIR3DL1*, both linear and non-linear kernel methods were used to predict the strength of interaction between the KIR3DL1 allotypes and any HLA class I allotype. First, we tested the prediction of binder/non-binder category, based on the information from all KIR3DL1 allotypes (see methods for training/test division) using a max-margin linear formalism mixing binary prediction and regression denoted MLVO ^24^ and a standard SVM. Note that more advanced algorithms, such as XGBoost and neural networks, were also tested, but given the limited amount of data their test accuracy was lower. The accuracy of linear classifiers suggests that the contribution of each residue to the log binding affinity is additive.

Disease association studies attempting to link the KIR3DL1/Bw4 interaction with clinical outcomes typically have only considered the presence or absence of the Bw4 motif or the presence of an Ile or Thr at position within this motif to affect the interaction with KIR3DL1. However, consistent with structural studies, the results above suggest that much larger regions of HLA class I are involved in binding to KIR3DL1. To assess this in more depth, the performance of each model was evaluated using four different options for the inclusion of amino acid residue positions (see methods for the definition of each region): 1) All positions (exon 2 and 3 encoding the *α*1 and *α*2 domains respectively), 2) all positions except connecting loops, 3) only positions on the alpha helices, and 4) only the six positions that contribute to the Bw4 motif itself. Since most Bw4 binding motifs are in the HLA-A and B genes^5^, we also evaluated the models with and without the inclusion of HLA-C (ABC and AB). While traditionally dimension reduction for categorical variables is performed using MCA, comparison of PCA and MCA results showed a preference for a PCA. Similar performance was obtained for MLVO and SVM, with the top AUC results obtained for the MLVO (Supplemental Figure 5).

When comparing different types of input, overall, there was no significant difference between the prediction accuracy on the test set for binder/non-binder observed with HLA-C excluded or included. However, a significant difference was observed when the six positions that contribute to the Bw4 motif were used versus when other positions were included (ANOVA p<0.05, Tukey test p<0.05, Figure 4A). We further used the only helices positions and with HLA-C, which was the minimal model with the top AUC (similar to everything but the loops).

**Figure 4:**
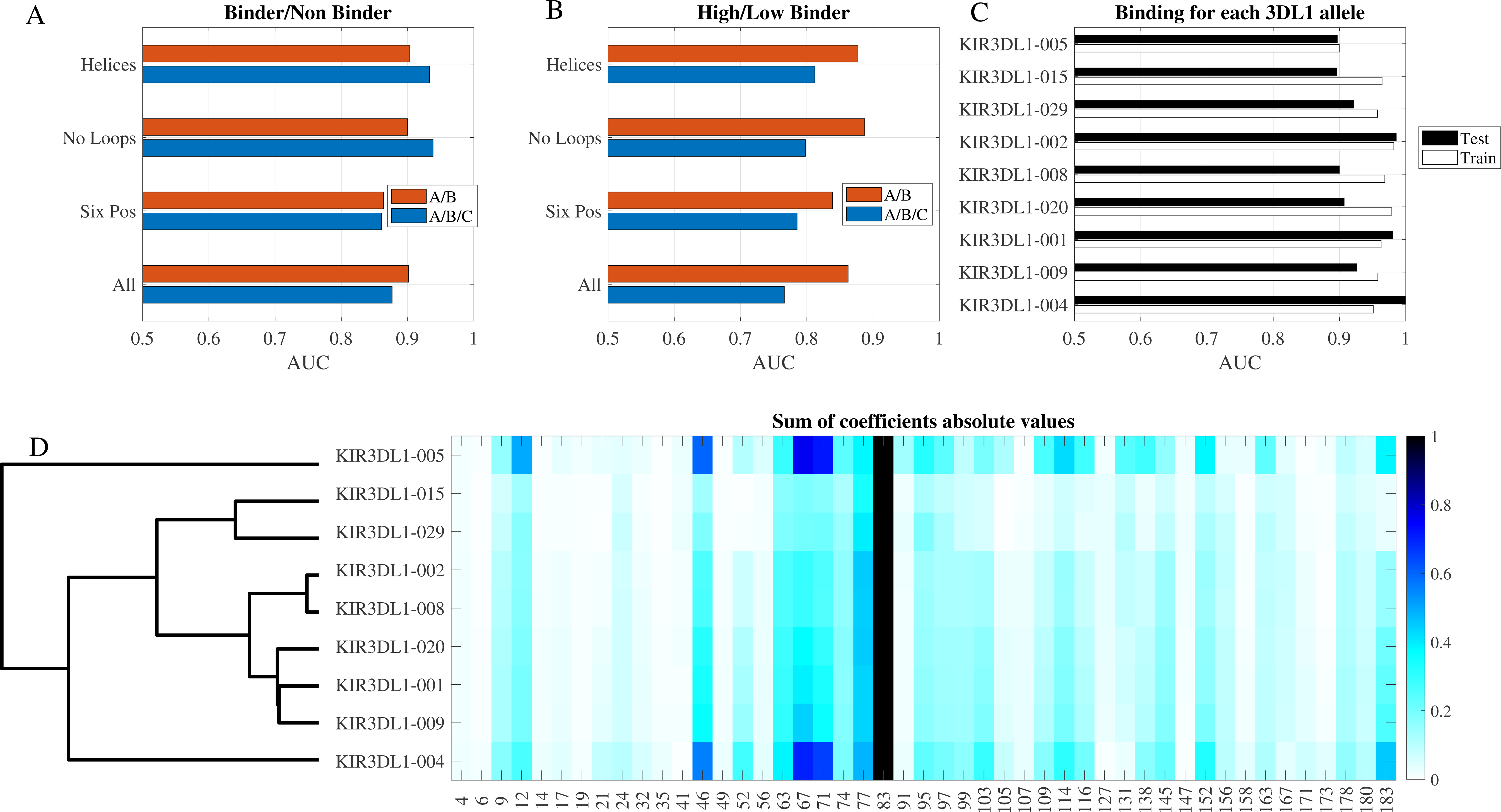
Machine learning results. **A**. Test AUC of classification into binders and non-binders, based on the clusters in Figure 3. All KIR3DL1 allotypes were used simultaneously. Each HLA allotype was either in the training or test set for all allotypes. We used different sets of positions in the HLA molecule for the classification and used either all loci or only the HLA-A and HLA-B loci. **B.** Same as A but for the classification between the high and low binder clusters. **C.** AUC of classifier per KIR3DL1 allotype using the no-loops and all HLA loci option. All AUC results in plots A-C are fivefold cross validation. **D.** Sum of absolute value of coefficients normalized by min-max (maximal value is 1, minimal value is 0). Each row is a classifier, and each column is a position. Only variable positions are marked. The allotypes are clustered and ordered according to their clustering (dendrogram to the left).

A similar analysis was performed to predict the high/low-binder category, using a linear classifier. In the high/low-binder classification, removing HLA-C from the analysis significantly reduced the performance (Figure 4B, ANOVA p<0.01, Tukey test <0.05 on all combinations of groups without HLA-C versus groups with HLA-C). This is expected from Figure 1, where many of the low-level binders are HLA-C allotypes. However, even when HLA-C was removed, a partial classification can still be obtained, since even within the HLA-A and -B allotypes there are differences between high and low-binders. The helices positions had the highest accuracy again and were again chosen for the remaining of the analysis.

We used the helical regions to predict HLA-I binders vs non-binders for each KIR3DL1 allotype separately. Interestingly, a clear difference was found between allotypes with KIR3DL1*005 performing worse, and other KIR allotypes like *002 having a test set AUC of almost 1 (Figure 4C), all based on the same number of observations (p<0.01 Tukey test of KIR3DL1*005 versus others). The linear classifier can be used to estimate the contribution of each position to the binding prediction (Figure 4D). The total contribution of each position was normalized to 0, and the average contribution of amino acids in each position is color-coded in Figure 4D. The divergent, broader binding pattern of KIR3DL1*005 likely drove its suboptimal performance in the model although KIR3DL1*004 performed better despite also having a divergent binding pattern. Beyond that, as expected, the main variance between positions was in residues 77-90, with some contribution from residues 67-71, mainly in KIR3DL1*005 and *004. Again, as expected from the different binding pattern (Figure 3), the classifiers are most different for KIR3DL1*004 and *005.

We computed the binder/non-binder and the high versus low binding scores for all KIR3DL1 allotypes and for 8,000 HLA A, B and C allotypes (Figure 5). Non-binders are mostly HLA-A allotypes, while the high binders are comprised of both HLA-A and -B allotypes (Figure 5A). The low binders are mostly HLA-C allotypes. When looking at the amino acid composition, binding corresponds with the I80 and T80-containing Bw4 motifs (Figure 5B and the presence of R83 in Figure 5C), while non-Bw4/C1 (N80/G83) and C2 (K80/G83) molecules sit in the middle range and the T80/G83 HLA-A allotypes did not bind at all. Analysis of position 66 (Figure 5D) did not reveal any single amino acid associated with high vs low binding despite variation this position significantly contributing to this distinction in the model. Some HLA-A allotypes with I80/R83 were non-binders (e.g. HLA-A*25:01), yet while other HLA-A along with HLA-B allotypes with I80/R83 are regarded as high binders. Overall, however, HLA-B allotypes tend to have higher predicted binding probability than HLA-A allotypes (Figure 5). Therefore, beyond the broad distinction between the main KIR ligands, the current analysis gives a much more graduated representation of KIR3DL1 and HLA-I binding.

**Figure 5:**
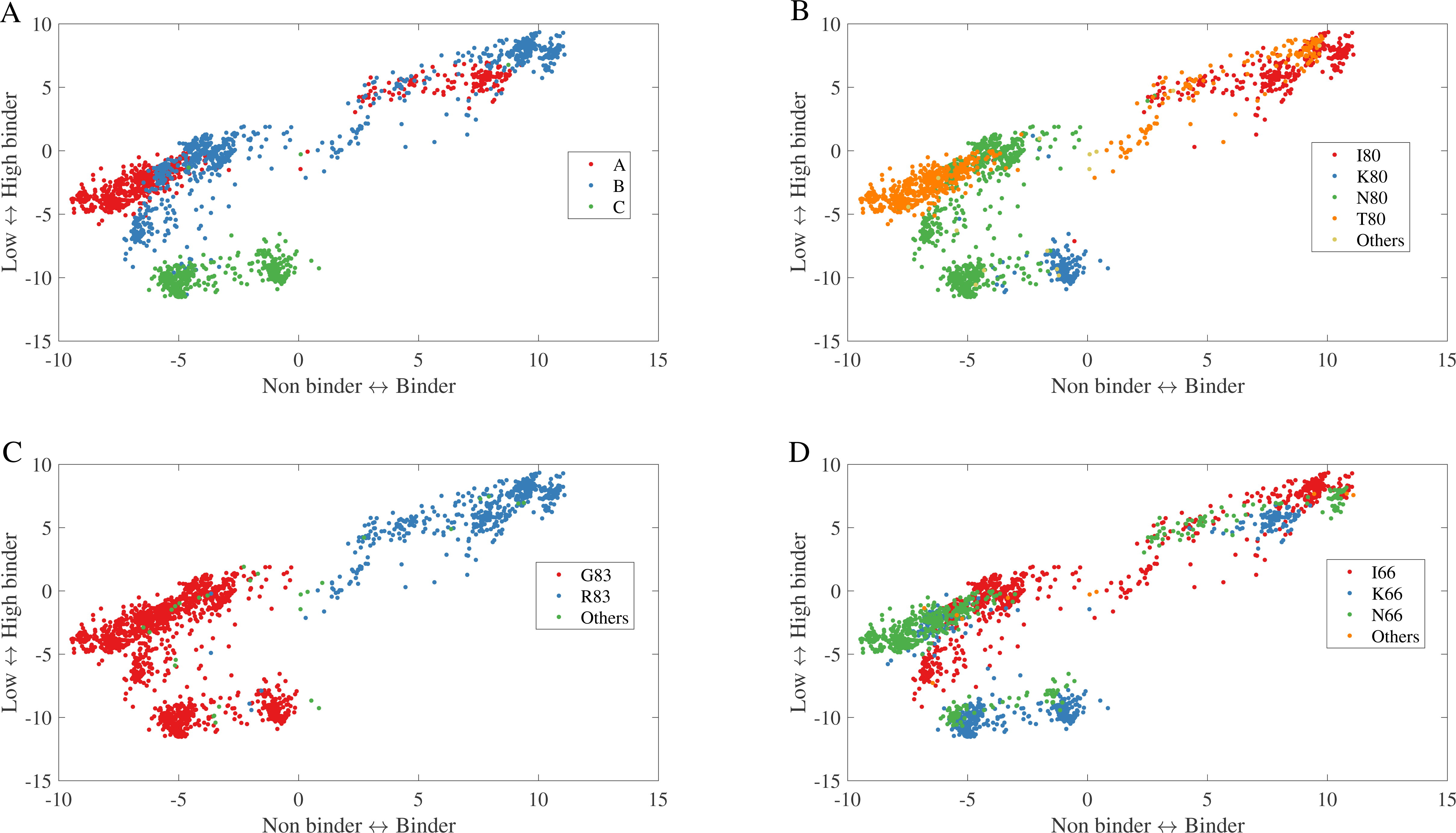
Projection of predictions. Prediction was performed for all HLA allotypes of the binder/ non binder classifier (x axis) and high-low binding classification (y axis). Allotypes are marked by their gene (A) and colored by their position 80 (B), 83 (C) and 66 (D) amino acids. Rare combinations were removed. One can clearly see the distinction between A/B and C (high vs low binders), C1 vs C1 (C binder vs non binder) and the Bw4 vs Bw6. However, beyond that, there is a large variability between HLA allotypes in binding score.

To test the accuracy of the predictor on unseen HLA allotypes, we used previously published data examining the recognition of wild type and mutant HLA by KIR3DL1^+^ NK cells. In these studies, HLA-I molecules were mutated at specific residues and transfected into 721.221 (221) cells, which lack endogenous HLA-A and -B allotypes. Purified NK cells from healthy blood donors typed for KIR3DL1 were then incubated with HLA-expressing 221 target cells and degranulation measured after five hours by flow cytometry. The mutations examined included residues comparing the recognition of HLA-B*57:01 and -B*13:01 (residue 145), HLA-B*57:01 and -A*24:02 (residues 144, 151, 116, 113/114/116, 95/97), and HLA-A*24:02 and -A*25:02 (residues 90, 149, 152), and the replacement of KIR3DL1 contact residues on HLA-A*24:02 and -B*57:01 with alanine (residues 16, 17, 18, 72, 76, 79, 80, 83, 84, 89, 142, 145, 146, 149, 150 and 151) ^10^ as well as our unpublished data examining residues 62 and 109, and comparing HLA-B*27:04/05/06 (manuscript in preparation).

We computed an expected HLA-KIR3DL1 binding affinity for each mutant HLA-I based on its amino acid sequence. Since individuals can express more than one KIR3DL1 allele, we computed the expected binding as the average binding rate for the KIR3DL1 alleles in an individual. To control for variability across HLA mutants and experiments, the observed degranulation of KIR3DL1^+^ NK cells was first normalized to their response towards the HLA-deficient 221 parental cell line in each assay (representative of the maximal degranulation). When this value was correlated with the computed binding affinity, a positive correlation was observed for all KIR3DL1 allotypes with an average of 0.4 (Figure 6A-B). Although the variable sample sizes here contribute to the differences in correlation, a higher accuracy is obtained for KIR3DL1 homozygotes (where the KIR allele is known).

**Figure 6.**
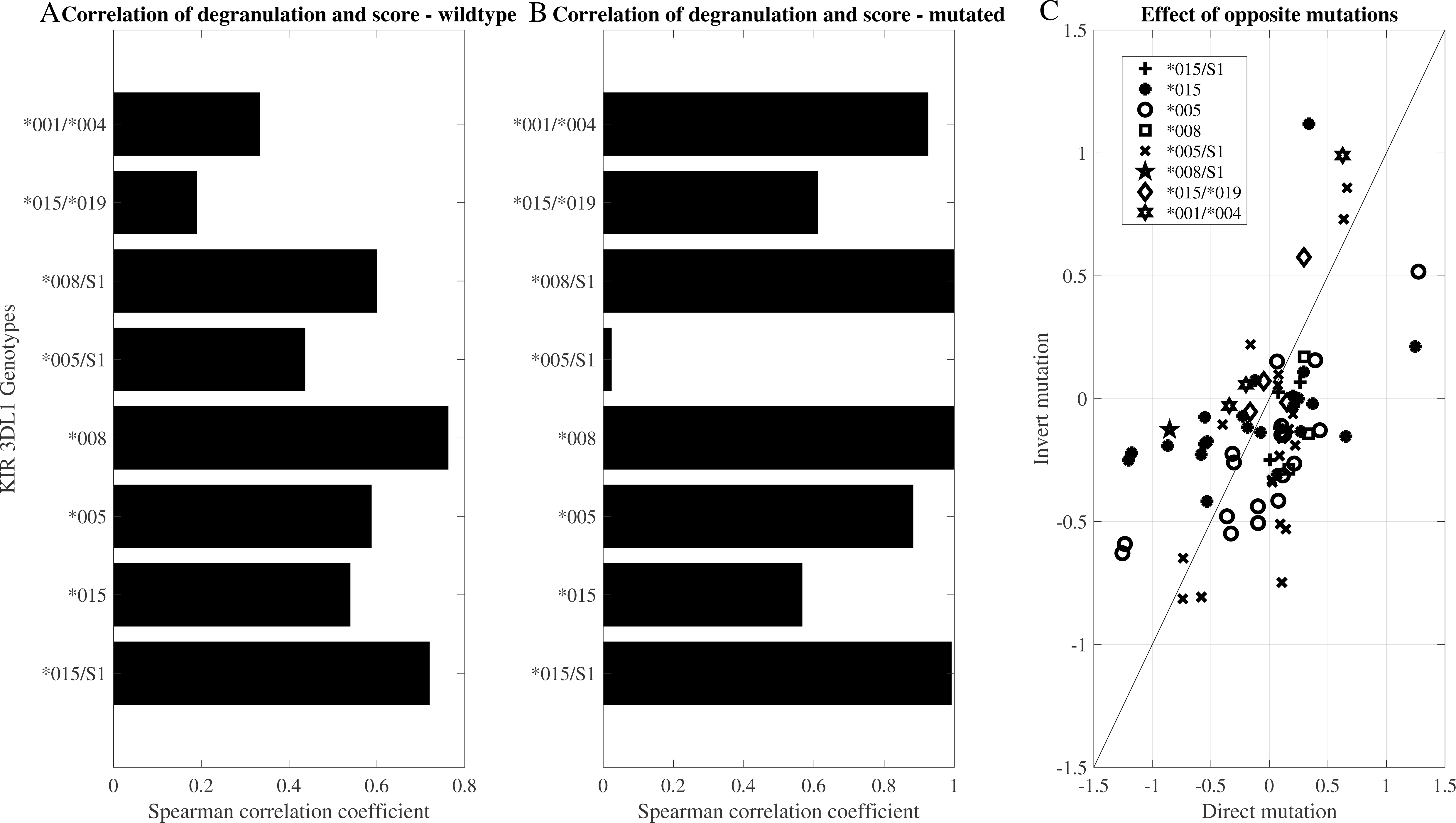
**A.** Correlation between prediction based on tetramer activation and observed degranulation of primary KIR3DL1+ NK cells towards transfected target cells. For heterozygous KIR3DL1+ NK cells, we computed the average predicted binding for the two 3DL1 alleles. **B**. Same correlation as in the left plot for mutated alleles. **C.** Comparison of the effect of opposite mutations on degranulation. Each different symbol is a cell line. Each point represents the change in binding affinity in a mutation (e.g. Q->K in position 144) and minus the effect of the opposite mutation (K->Q in position 144 on a different allele background). If the contribution of different amino acids was additive, we would expect all points to be on the diagonal.

The model used here presumes a linear contribution of each amino acid to the log of the binding affinity. As such, mutation of a given amino acid would be expected to have an opposite log ratio change upon mutation in the reverse direction. To test for this, we normalized the degranulation of KIR3DL1^+^ NK cells towards each mutant HLA-I with the response seen against the wild type HLA-I allotype. We then computed the log ratio of the change in degranulation and compared it to the log of the degranulation change for the opposite mutation. For example, assume an HLA-A*24:02 allotype with a Q->K mutation at position 144, and an HLA-B*57:01 K->Q mutation. One would expect the log ratio of the degranulation to be the inverse. However, in this case, both mutations decrease or do not change the degree of degranulation.

To test how consistent the linearity assumption is, we correlated the log ratio of the degranulation of KIR3DL1^+^ NK cells towards a given HLA-I mutant to the log ratio of the opposite mutation (Figure 6C). When assessed by KIR3DL1 allotype, the responses of KIR3DL1*005^+^ NK cells to reciprocal HLA-I mutations correlated more closely to one another than the response of KIR3DL1*015^+^ NK cells to such mutations. Given that the binding affinity of KIR3DL1*015 displayed greater variability across HLA-I than KIR3DL1*005 (Figure 1), the sensitivity of KIR3DL1*015 to these reciprocal mutations is likely affected by its capacity to bind HLA-I of different allotypes. In contrast, the broader recognition of HLA-Bw4 allotypes by KIR3DL1*005, as well as its greater peptide tolerance ^7^ may allow for more reflective impacts by individual HLA-I mutations. These results highlight both the prediction strength of the general model developed here as well as some of its limitations.

### Webtool

A webtool has been developed (https://kir-hla.math.biu.ac.il) that allows input of either an HLA-A, - B, or -C allele name or the amino acid sequence of an allotype (including novel or hypothetical alleles) and predicts their likely binding to KIR3DL1 allotypes. The output from 11 models is provided as described in the methods along with an indication of how this allele performs relative to the full list of HLA-A, -B, and -C alleles evaluated. The matrix of beta values for all 11 models is available for download as well as the amino-acid encoding for 15,474 HLA-A, -B, -C alleles (as of 2023-04-01). (Supplemental figures 1-4)

## 4 Discussion

KIR3DL1 binding to HLA at an individual allotype level has been implicated as a determinant in numerous clinical contexts: the immune response to cancer and transplants^29,30^, the response to viruses^31,32^ and even the development of neurological disorders^33,34^. Beyond those, KIR3DL1-HLA-Bw4 interaction ^29^ is implicated in the success of immunotherapy. Models that assume their binding is solely based on the Bw4 epitope lose much of the complexity of HLA-I interaction with KIR3DL1. For example, the simplistic presence of I80/R83 would predict KIR3DL1 binding to HLA-A*25:01, which is not the case. While there are experimental measures of KIR3DL1 binding to multiple HLA allotypes, the probability of a patient having experimental measures of all their HLA-I and KIR3DL1 alleles is below 30% even for the best covered populations. To address this gap in experimental coverage we have characterized the HLA-KIR3DL1 binding and developed a machine learning tool to predict overall KIR3DL1 binding to HLA for any HLA class I allotype and binding to nine specific KIR3DL1 allotypes.

Three important observations arise from both the prediction algorithm and directly from the binding measurements: A) there are three distinct levels of binding between KIR3DL1 and HLA-I allotypes (high-binding, low-binding and non-binding); B) the binding is not only determined by position 77-83 (which defines the Bw4 motif), but there is a contribution to the binding affinity (and thus binding prediction probability) from remote positions (Figure 7). Thus, when predicting the KIR3DL1-HLA binding, the full surface of the HLA should be considered; and C) while most KIR3DL1 allotypes have similar binding patterns and as such a similar predictor, KIR3DL1*004 and *005 are different from all other allotypes. Specifically, these two allotypes tend to bind HLA class I that diverge beyond the classical Bw4 motif much more than others, with more contribution to the binding from regions beyond positions 77-83. Conversely, allotypes KIR3DL1*015 and *029 are more consistent with binding the classical Bw4 motif. These predications are consistent with functional NK cell studies showing KIR3DL1*005^+^ NK cells to have broader HLA binding capacity and greater peptide tolerance^7,19^.

**Figure 7.**
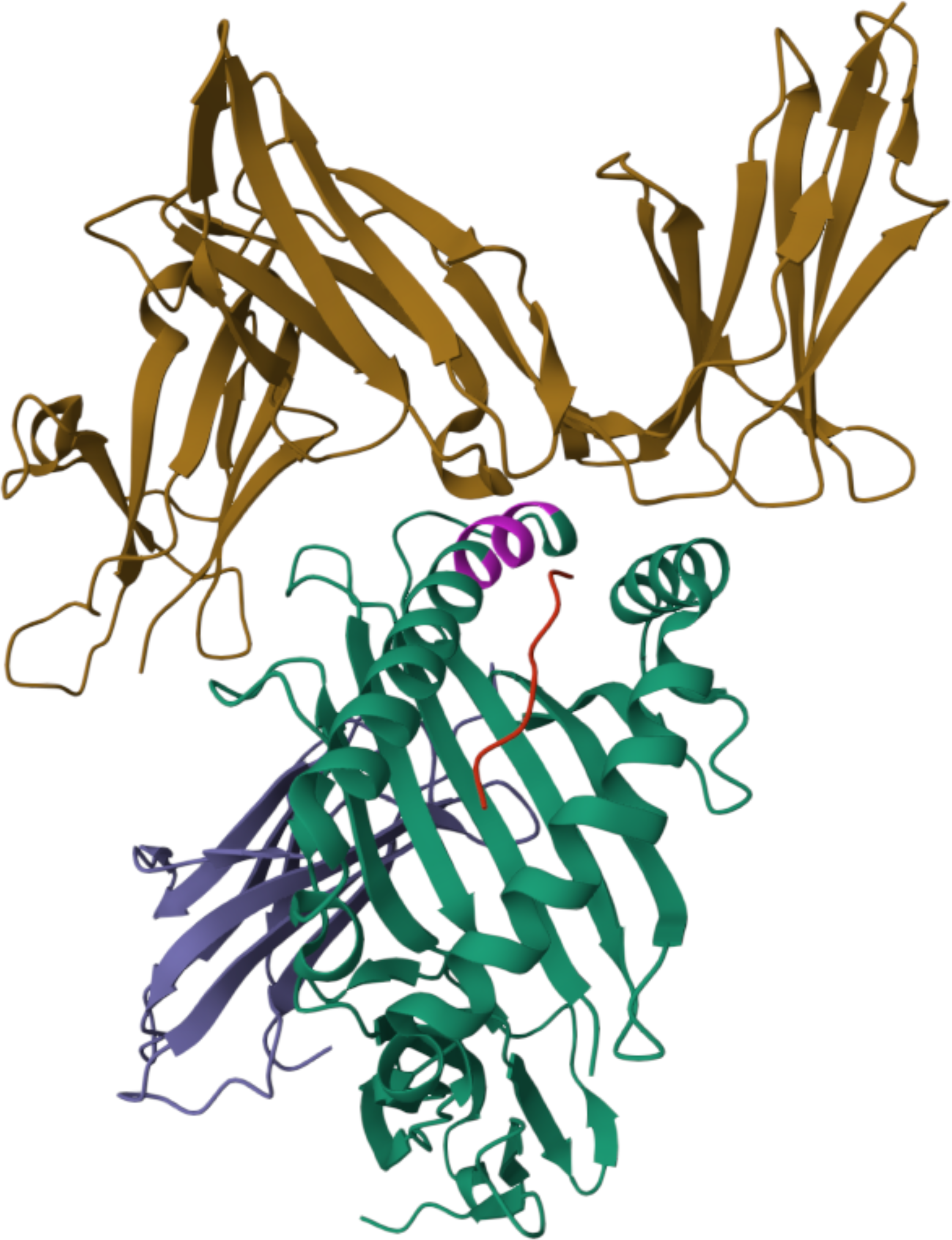
Structure of 3DL1*001 (brown) and HLA-A*24:02 (green) with RYPLTFGW peptide(red) (PDB accession 7K80). The Bw4 region is highlighted in pink. The Bw4 region is directly interacting with the 3DL1. However, other positions, distant from Bw4, contact the 3DL1 molecule.

The data here showed that a simple linear predictor obtains a high AUC on most KIR3DL1 allotypes, except for KIR3DL1*005, which has a slightly lower AUC. Moreover, the coefficients of the regressor are consistent in positions 77 to 83 with detailed mutagenesis experiments^7,8,19,30^, but the classifier adds information from other positions to improve the AUC.

Although the model correctly assigns HLA-A*25:01 as a non-binder (Supplemental Figure 1), despite the presence Ile at position 80, it is worth noting there are still outliers such as HLA-B*13:01 which is classified as a binder in the overall KIR3DL1 model despite displaying no interaction with KIR3DL1 in the binding assays and failing to mediate efficient KIR3DL1^+^ NK cell inhibition in functional assays^10^, Nevertheless, the individual models for the nine KIR3DL1 allotypes reflected the training data of low binding for most HLA alleles but still detected slightly higher binding for KIR3DL1*004 and KIR3DL1*005 (Supplemental Figure 2).

Prediction of binding between KIR and HLA presents inherent challenges due to the influential role of peptides in modulating binding affinities. For example, functionally distinct responses to KIR3DL1 engagement between HLA-B*57:03 vs HLA-B*57:01, which differ only at positions 114 and 116 in the floor of the peptide binding groove, have been shown to be due to conformational differences in the peptides associated with each allotype^9^. Developing models that can accommodate these complex peptide sequence/confirmation effects will be challenging. Moreover for KIR2D– HLA-C interactions, both activating and inhibitory receptors, have recently been shown to exhibit a wide range of peptide specificity ^26^ offering prospects for future KIR/HLA/peptide prediction. Further complicating the prediction of sequence-based binding prediction is the recent discovery of plasticity in the binding where KIR2DL recognized the C1 allotypes HLA-C*03:04 and HLA-C*07:02 peptide complexes using different molecular contact points^27^. Future binding prediction models will need to account for this plasticity in binding since these data challenge the assumption that a specific residue position has the same involvement across allotypes.

Despite these challenges, we see a role in applying successively more refined models of binding between KIR and HLA to clinical applications such as transplantation and other cellular therapies. The potential for a graded system of high/low/non-binding for the KIR3DL1 and HLA-B interaction in the context of hematopoietic cell transplantation for acute myelogenous leukemia (AML) was shown by Boudreau et al^31^. Their approach considers the expression of KIR3DL1*004 as a “null” allele which is not considered in the binding assays used in the machine learning model we have developed. This model did not validate on an independent cohort^32^. In contrast, our approach considers the important and well-established capability of the Bw4 epitope expressed on HLA-A to bind KIR3DL1 and inhibit NK cells. The additional resolution on both the HLA and KIR allotypes provided by our approach has the potential to offer more specific re-analysis of the transplant cohorts.

This predictor has wide potential clinical applications based on known associations between HLA and KIR3DL1 in transplantation, cancer immunotherapy and disease association studies. As more experimental data becomes available, we envision improving these predictions, may ultimately lead to improved clinical decision making as well as better elucidation of this important immune interaction.

## 7 Author Contributions

MM designed the study. MM and YL wrote the manuscript. YL performed machine learning. MM and YL analysed results. PS contributed experimental data and text and reviwed the manuscript. AB, PP, JV, JR generated binding data and reviewed the manuscript.

## 9 Data Availability Statement

The datasets analyzed for this study can be found in the Supplementary Materials file.

---

**Supplemental Figure 1.** A screenshot of the webtool (https://kir-hla.math.biu.ac.il/) demonstrating the prediction based on an input of an HLA allotype (e.g. A*25:01).

**Supplemental Figure 2.** A screenshot of the webtool (https://kir-hla.math.biu.ac.il/) demonstrating the prediction based on an input of an HLA allotype (e.g. B*13:01).

**Supplemental Figure 3.** A screenshot of the webtool (https://kir-hla.math.biu.ac.il/) demonstrating the prediction based on an input of the amino acid sequence encoding the *α*1 and *α*2 domains of an HLA Class I allotype

**Supplemental Figure 4.** A screenshot of the webtool (https://kir-hla.math.biu.ac.il/) demonstrating the prediction based on an input of an HLA allotype (e.g. A*24:02).

**Supplemental Figure 5.** Comparison of the distribution of AUC values (y axis for different dimension reduction algorithms (MCA vs PCA) and linear predictors MLVO vs SVM. Non-linear predictors, such as XGBoost and fully connected neural networks had worse performance. Each dot is a different train/internal validation division.

