## Supplemental Figure 3 for "Prediction of KIR3DL1/Human Leukocyte Antigen binding"

### KIR HLA allele binding

Please enter an amino acid sequence **or** its HLA allele:

EQRRAYLEGTCVEWLRRYLENGKETLQRAE calculate

**C*01:02**
**Score for binding: -11.081**

The blue bars represent the scores for each 3LD1 allele. The red bar represents binding to 3LD1 in general (model trained on all alleles simultaneously). The green bar represents a model for the difference between low and high binders. The interpretation of the bars should be first to check for binding, then, if an HLA allele is a binder, look for high vs low binding.
Alleles with positive values are typically binders.

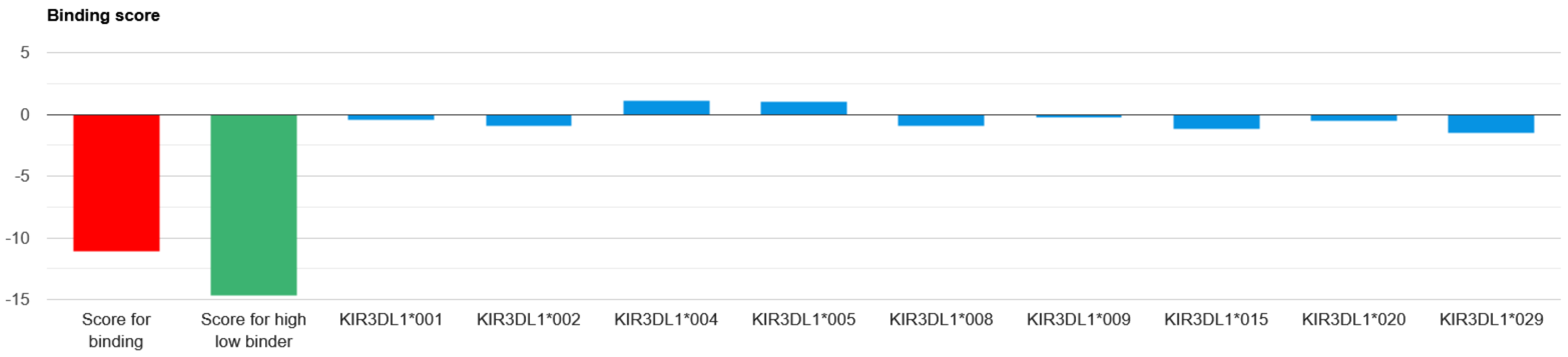


The lower plot represents the percentile of the score for each allele or over all alleles as above. The empty bars represent the threshold percentile for any 3LD1, alleles with percentiles above the thresholds are typically binders.

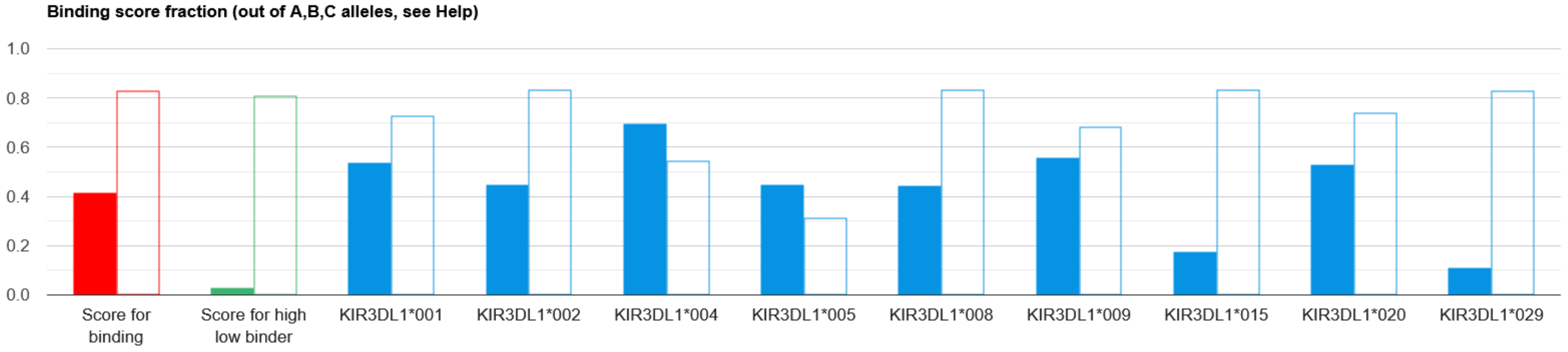
