## Supplementary figures and images for "Prediction of KIR3DL1/Human Leukocyte Antigen binding"

### Supplemental Figure 5

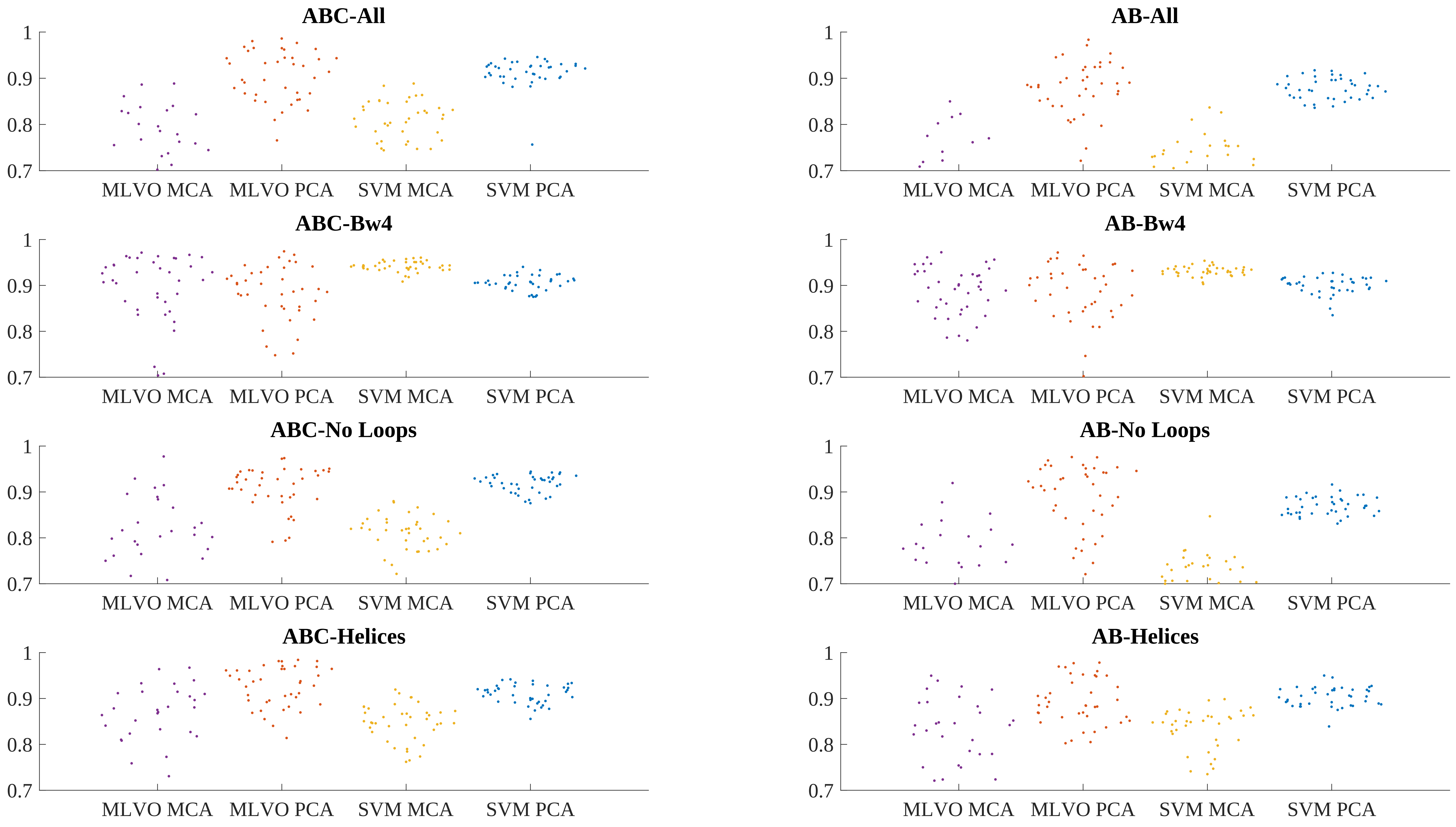
